# Priority effects influence bacterial community assembly on temperate deciduous tree leaves

**DOI:** 10.64898/2026.10.05.756808

**Authors:** Zihui Wang, Vahid J. Javaran, Jocelyn Lauzon, Jean-Baptiste Leducq, Steven W. Kembel

**Author notes:** co-first authors (Zihui Wang, Vahid J. Javaran).

## Abstract

Microbial community assembly on leaves (the phyllosphere) is influenced by deterministic, stochastic, and historical processes. However, the importance of historical processes such as priority effects, where the timing and order of colonization influence community assembly, has not been extensively investigated in field-based phyllosphere studies. We designed a field experiment in a temperate forest in Quebec where we inoculated five strains of *Methylobacterium* onto the leaves of red maple and American beech trees, either early or late during the growing season. We quantified microbial community structure throughout the season using metabarcoding of a *Methylobacterium*-targeting *rpoB* primer as well as a 16S rRNA primer targeting the entire bacterial community. We show that the effects of inoculation on bacterial communities depend on inoculation timing: early-season inoculation led to strain-specific shifts in community composition, while late-season inoculation had no significant effect. Although the effect of early-season inoculation weakened over time, early- and late-season inoculation still led to different community compositions at the end of the growing season. Our study demonstrates that priority effects contribute to the assembly of phyllosphere bacterial communities under field conditions, with arrival timing and strain identity shaping community trajectories. Considering historical processes will thus be crucial for successful microbiome management.

## Introduction

Microorganisms living on leaves play critical roles in supporting plant health, protecting plants against pathogens, and in influencing plant and ecosystem productivity and function [1,2]. Because of this, there is increasing interest in manipulating or engineering the leaf (phyllosphere) microbiome to optimize plant health and ecosystem services, particularly in agriculture and forestry [3,4]. Promising approaches including the use of bioinoculants or synthetic communities in these contexts have often led to inconsistent results, as introduced strains may fail to establish or to have lasting impacts on the plant host in field conditions [5,6]. Successful microbiome engineering will thus require a fundamental understanding of the ecological processes governing the assembly of plant-associated microbial communities [7]. While numerous deterministic and stochastic processes have been shown to influence phyllosphere community assembly, less is known about the role of historical contingency for community assembly. In particular, the impact of priority effects [8], where the timing and order of microbial taxa arrival impacts community assembly, has not been widely considered in models of plant microbiome assembly in the field conditions.

The assembly of phyllosphere bacterial communities initiates as emerging leaves are colonized by microbes from the air, soil, seed, and other sources, followed by a succession shaped by both stochastic and deterministic ecological processes [9–11]. Phyllosphere community assembly has been shown to be influenced by host plant selection, environmental filtering, and continuous colonization from the surrounding metacommunity [12–14]. Understanding the relative importance of these processes is key for developing effective microbiome engineering strategies. For example, if host filtering strongly influences community assembly, microbiome engineering may be improved by breeding plant genotypes whose traits favour the recruitment and persistence of desired microbes. In contrast, if assembly is strongly influenced by stochastic dispersal, success may depend more on applying target microbes at sufficient densities and increasing their opportunities for colonization [15]. Field-based studies have reported conflicting evidence on the relative importance of these assembly processes, as their importance can vary greatly in both time and space, and among study systems [13,16,17].

Priority effects represent an important but understudied community assembly process in field settings, where the timing and order of arrival of colonists influence community assembly through niche pre-emption or niche modification [18,19]. There is ample theoretical support and empirical evidence for the importance of priority effects for community assembly in diverse habitat types [8]. For phyllosphere communities in field settings, the evidence for priority effects is much more limited compared with lab- and greenhouse-based studies [20,21], and their relative importance compared with deterministic and stochastic assembly processes has not been widely quantified [11]. We expect that the importance of priority effects should be reduced when there is strong selection driving succession towards a climax community state, or when high rates of immigration and colonization homogenize community composition [19,22,23] and lessen the impact of early colonization.

In the temperate forest phyllosphere, priority effects may influence microbial community succession on leaves throughout the growing season. On deciduous trees whose leaf emergence is relatively synchronous early in the season, newly emerged leaves represent a habitat with relatively low microbial biomass and abundant niche space. These characteristics create an important opportunity for early leaf colonists to establish successfully as dominant taxa [9,11] and to potentially have strong impacts on subsequent community assembly [8]. As the growing season progresses and leaves and their communities mature, microbial biomass accumulates and niche space saturates. This is paralleled by potential increases in the strength of biotic and abiotic selection on community assembly, or in the rate of taxa dispersal from the surrounding environment [12,24,25]. As a result of these interacting processes, we would expect late leaf colonists to face greater resistance to establish as dominant taxa and, when dispersal is strong, to observe greater homogenising impacts on communities driven by mass effects. This homogenization would eventually lead communities to converge towards a relatively homogenous climax compositional state, with a reduced impact of late colonization by specific taxa [20,26].

In this study, we used a field inoculation experiment to quantify the importance of priority effects on phyllosphere community assembly on the leaves of two tree species, red maple and American beech, in a temperate deciduous forest. We inoculated leaves with five distinct strains of *Methylobacterium,* a key phyllosphere genus [10], at two successional stages during the growing season (early and late season) and quantified microbial community structure on inoculated leaves and uninoculated control leaves two weeks after inoculation and at the end of the growing season. We predicted that inoculation would increase the relative abundance of taxa matching the inoculated strains and affect the abundance of other bacterial taxa. We then tested two hypotheses regarding the role of priority effects in community assembly. First, inoculation would affect bacterial community assembly differently when strains were introduced early versus late in the growing season, with inoculation and strain identity having a stronger effect when inoculating newly emerged leaves compared with mature leaves. Second, the importance of inoculation in shaping community assembly should decrease over the course of the growing season, as host and environmental filtering, microbial dispersal and seasonal succession become increasingly important.

## Methods

### 1. Study site

We established a field inoculation experiment to quantify the importance of priority effects on phyllosphere bacterial community assembly in forests at the Station de biologie des Laurentides in Quebec, Canada (45.99°N, 74.01°W). The station is located in the lower Laurentian region of the Canadian Shield and comprises approximately 16.4 km^2^ of hilly terrain containing temperate forests, lakes, wetlands, and streams. This protected area supports a diverse regional biota, with a canopy mainly dominated by white birch (*Betula papyrifera*), poplar (*Populus spp.*), red maple (*Acer spp.*), and American beech (*Fagus grandifolia*). Previous work at this site showed that leaf bacterial communities experience seasonal change, host filtering, and repeated microbial immigration from the surrounding landscape [10,14,27].

### 2. Selection of strains

In 2018, we conducted a survey of leaf-associated *Methylobacterium* strains at the Station de biologie des Laurentides using both metabarcoding- and isolation-based approaches [10]. We selected five strains corresponding to the most abundant and representative subclades of the *Methylobacterium* genus in the field: strains J-077, E-041, J-118, E-012, and E-025 (table S1). In our survey, these strains displayed different ecological attributes. For example, J-077 and E-012 were more abundant late in the growing season, while J-118 and E-025 were more abundant during early season [10]. E-041 was more closely associated with American beech (*Fagus grandifolia*), while J-077 and J-118 were more closely associated with red maple (*Acer rubrum*) (table S1).

The selected strains were previously isolated and preserved in 25% glycerol at -80°C. Their identity and purity were confirmed by whole-genome sequencing before the experiment (NCBI BioProject PRJNA730554). To prepare strains for inoculation, each strain was revived and cultured in ten Petri dishes on a R2A agar medium supplemented with 3.5 mM succinate at 23°C for two weeks.

After incubation, dishes were inspected to confirm healthy colony growth with no visible contamination. We then added 10 mL of sterile 10 mM MgCl₂ solution to each dish to gently resuspend bacterial cells, and suspensions were diluted with sterile 10 mM MgCl₂ solution to a standardized OD₆₀₀ of 0.015 [28,29]. In total, we generated 5 distinct bacterial strain inocula and a sterile 10 mM MgCl₂ solution as the control treatment.

### 3. Experimental design (inoculation and sampling)

In May 2022, shortly after initial spring leaf emergence, we established two 20×20m study plots, approximately 1.2 km apart, in mature forests at the study site. Within each plot, we randomly selected six trees belonging to each of the two focal host plant species, American beech (*Fagus grandifolia*) and red maple (*Acer rubrum*), for a total of 24 trees across species and plots (figure 1). The two species were selected because they are dominant deciduous trees at the study site, differ in leaf and ecological traits and could be readily accessed and monitored throughout the season. Focal trees were spatially distinct, with non-overlapping crowns and a diameter at breast height ranging from 2 cm to 10 cm. Each tree had multiple naturally occurring branch segments in their subcanopy (2-3 m height). For each tree, we identified six spatially separated branch segments, each bearing 10-20 healthy individual leaves with no obvious signs of disease or herbivory damages. We randomly assigned these six branch segments to one of six treatments: inoculation with one of the five *Methylobacterium* strains or a control treatment. Each branch segment and its assigned treatment was marked with aluminum tags to permit tracking of leaves throughout the growing season.

**Figure 1.**
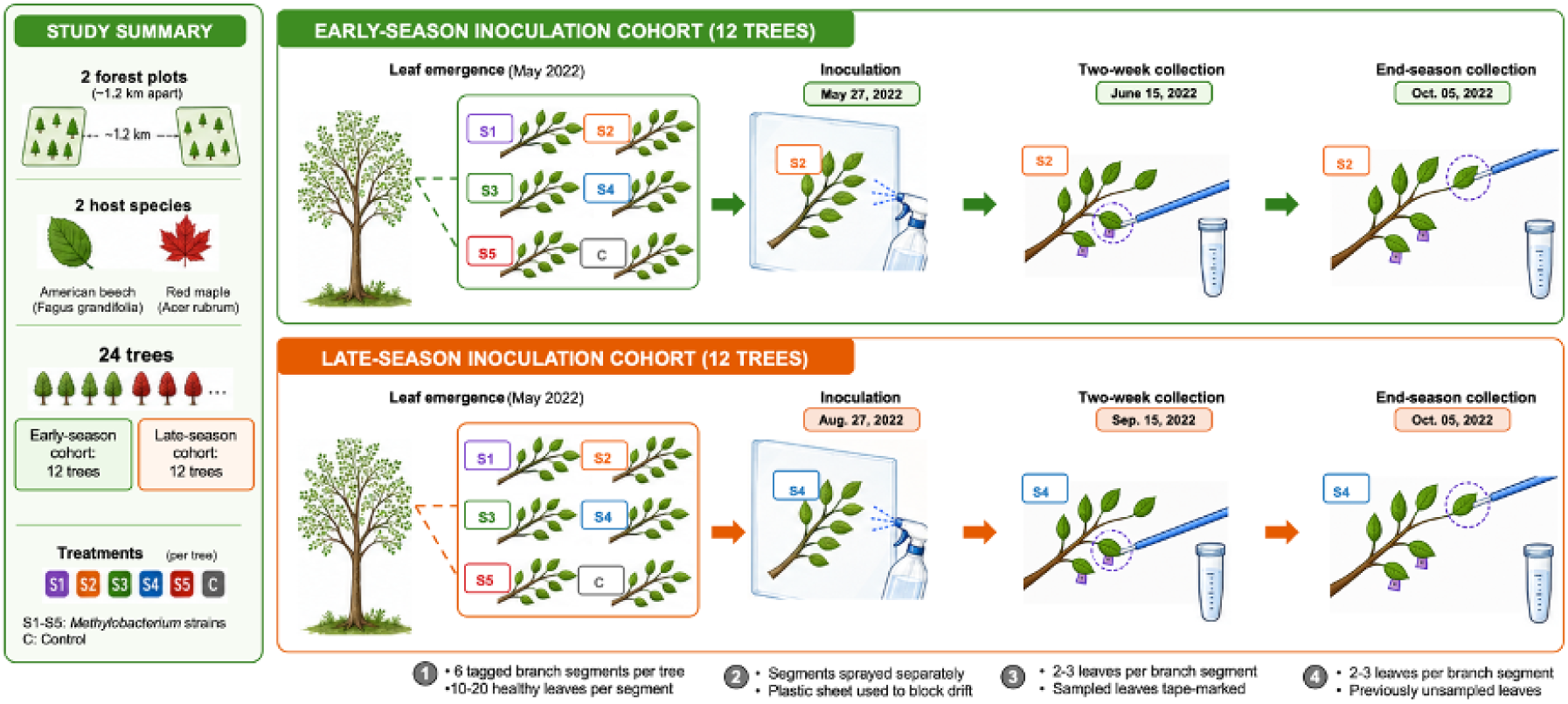
Experimental design of leaf microbial inoculation and sampling. The experiment was conducted in two forest plots using two focal host plant species, American beech (*Fagus grandifolia*) and red maple *(Acer rubrum*). A total of 24 trees were divided into early- and late-season inoculation cohorts. Within each tree, six tagged branch segments were assigned to one of five *Methylobacterium* strain treatments (S1-S5) or to a control treatment. Branch segments were sprayed separately during inoculation, and microbial communities were sampled from leaves two weeks after inoculation (two-week collection) and again at the end of the growing season (end-season collection). Leaves sampled at the first collection were tape-marked, so the end-season collection targeted previously unsampled leaves on the same branch segment.

We randomly divided all the trees into two groups, each with 12 trees (3 individuals from each species in each plot) into either an early-season or a late-season inoculation cohort (figure 1). We applied the inoculation treatment to each tree by isolating each branch segment and using a handheld spray bottle to apply either *Methylobacterium* inoculum or sterile control liquid. During spraying, each branch segment was temporarily isolated from neighbouring branches using a plastic sheet. The nozzle was held close to the leaves, and the inoculum or control was applied evenly to both leaf surfaces until all leaves on the segment had been sprayed. Each branch received approximately 7-10 mL of inoculum or sterile solution. After treating each tree, we checked that treated branch segments were not in lateral contact and did not vertically overlap with one another, thereby minimizing the potential for inoculum transfer through physical contact or rainfall. Trees assigned to the early-season inoculation treatment were inoculated on May 27, 2022. Trees assigned to the late-season inoculation treatment were inoculated on August 27, 2022.

Microbial communities on inoculated leaves were sampled at two time points during the growing season, at two weeks post-inoculation (two-week collection), and at the end of the growing season (end-season collection). Two-week post-inoculation samples were collected on June 15th for early-season inoculated trees and on September 15th for late-season inoculated trees. End-season samples were collected on October 5th for both early-season and late-season inoculated trees, just before leaf senescence (figure 1).

Microbial community samples were collected using a sterile swab that was wetted with 0.3 mL of sterile phosphate-buffered saline (PBS) solution. We swabbed 2-3 leaves with a combination of horizontal, vertical, and diagonal movements throughout the upper and lower leaf surfaces. After swabbing, tips were deposited in a sterile 1.5mL tube and stored at 4°C for up to 48 hours prior to DNA extraction in the lab. The swabbed leaves were tape marked so that the following collections could sample microbial communities from previously unswabbed leaves. In summary, we collected a total of 288 microbial samples (24 trees x 6 branch inoculation treatments x 2 time points).

### 4. DNA extraction, metabarcoding sequencing and bioinformatics

We used DNeasy® 96 PowerSoil Pro HT Kit (Qiagen, Hilden, Germany) for DNA extraction. We first transferred the swab with a sterile forceps into the bead-beating tube (PowerBead Pro Tubes) and then added 800 µl of CD1 buffer. The tubes were horizontally vortexed at maximum speed for 10 min. We then centrifuged the beadbeating tubes at 15,000 x g for 1 minute and transferred the supernatant into new tubes. Further procedures followed the manufacturer’s protocol.

For each sample and for a PCR-negative control, we carried out PCR reactions to create sequencing libraries for two bacterial barcode genes: a host-DNA excluding 16S primer to quantify overall bacterial community composition (16S rRNA primers 799F-1115F [30]), and a *Methylobacterium*-targeting *rpoB* primer to permit additional resolution of individual *Methylobacterium* strains that was not possible with universal 16S rRNA primers (*rpoB* primers Met02-352F-Met02-1121R [10]). PCR and library preparation conditions followed previously described protocols [10]. The 16S rRNA and *rpoB* sequencing libraries were sequenced on an Illumina MiSeq at the UQAM CERMO-FC Genomics Platform. Paired-end 16S rRNA gene reads were processed in R with DADA2 v1.36.0 [31]. Primer sequences were removed [32,33] and reads were truncated to 260 (forward) and 185 (reverse) nucleotides before expected-error filtering. ASVs were inferred with pseudo-pooling, paired reads were merged, and chimeric sequences were removed using the consensus method. Taxonomy was assigned against SILVA SSU Ref NR99 v138.2 [34,35]. We removed 38 chloroplast and 34 mitochondrial ASVs and four samples with fewer than 1,000 reads (including the PCR-negative control sample), leaving 284 samples containing 5,898,121 reads and 8,101 ASVs.

*rpoB* amplicons were also processed using DADA2. Reads were quality-filtered and trimmed to remove the first 20 bases, retaining forward and reverse fragments of 225 and 165 bp, respectively. Forward and reverse reads were denoised separately using pseudo-pooling, paired reads were concatenated, and chimeras were removed with the consensus method. Genus-level taxonomy was assigned using the DADA2 naïve Bayesian classifier (minimum bootstrap confidence 50) against a custom full-length *rpoB* database containing 3482 *Alphaproteobacteria* and *Methylobacteriaceae* reference sequences [10]. Species-level annotations were obtained by exact matching of the forward and reverse fragments of each ASV against the *rpoB* species database using DADA2 “assignSpecies”. 1,417 ASVs and 610,088 reads assigned to *Methylobacterium* across 272 of 288 biological samples were retained for the following strain-level analyses.

### 5. Statistics and analyses

#### Rarefaction and sequencing-depth standardization

Rarefaction curves generated from the filtered 16S rRNA ASV table approached saturation at approximately 2,000–3,000 reads per sample (figure S1). Samples were randomly rarefied once to 3,448 reads per sample without replacement using vegan::rrarefy with a fixed seed. This retained 283 of 284 samples and produced a rarefied dataset containing 975,784 reads and 6,982 ASVs for diversity analyses. Because the rpoB dataset was used only to quantify and track the inoculated strains, rather than diversity analyses, rarefaction was not performed for this dataset.

#### Detection of inoculated Methylobacterium strains using rpoB marker

To assign *rpoB* ASVs to inoculated *Methylobacterium* strains, the forward and reverse fragments of each ASV were aligned separately by local pairwise alignment to full-length *rpoB* reference sequences for the five inoculated strains. Only rpoB ASVs assigned to *Methylobacterium* were retained for strain level analyses. An ASV was assigned to aa inoculated strain when it uniquely matched its reference at 100% nucleotide identity across at least 99% of its combined sequenced length. The relative abundance of each inoculated strain within *Methylobacterium* communities was calculated as the proportion of *Methylobacterium* reads assigned to its inoculant-matched *rpoB* ASVs; samples without *Methylobacterium* reads were excluded.

#### Differential-abundance analysis of 16S rRNA ASVs

Differential abundance between inoculated and control branches was tested using non-rarefied ASV counts. Early and late inoculations were analysed separately at two weeks post-inoculation and at the end of the season. Each sampling time was analysed across both host plant species and separately for red maple and American beech. ASVs detected in at least four samples and represented by at least 10 total reads were retained. Each strain was compared with controls using ANCOM-BC2 [36], LinDA [37], MaAsLin2 [38], ALDEx2 [39] and DESeq2 [40]. All differential-abundance comparisons accounted for the paired experimental design by comparing inoculated and control branches within the same host trees, using method-appropriate blocking or paired tests. Additional internal prevalence filtering was disabled, DESeq2 used positive-count size-factor estimation, and ANCOM-BC2 results were required to pass the pseudocount-sensitivity test. P-values were adjusted across ASVs within each strain contrast using the Benjamini–Hochberg procedure, recomputed for MaAsLin2 over the treatment coefficients alone; no further correction was applied across strains, sampling time or host scopes. Associations were retained when at least two methods agreed in direction and reached the same significance threshold. Support at (q<0.05) was classified as high confidence and support at (q<0.10) as moderate evidence. Effect estimates were not averaged because their scales differed among methods.

#### Effects of inoculation on bacterial community composition

We used Permutational Multivariate Analysis of Variance (PERMANOVA) to test the effect of inoculation on bacterial community composition All PERMANOVA analyses used Aitchison distances calculated from rarefied 16S rRNA ASV table after adding a pseudocount of one.

Distances were calculated using ‘vegdist’ and models were fitted using ‘adonis2’ in the vegan package (v2.6.10) [41] with 999 unrestricted permutations. In all models, host plant species, plot identity and their interaction were entered before the inoculation terms. Therefore, inoculation effects were measured after accounting for host and plot effects.

To test short-term inoculation effects (two weeks post inoculation), we conducted PERMANOVA tests separately for early- and late-season inoculation cohorts, as these samples were collected in different times in the season and the sampling time would otherwise confound with inoculation timing. Within each inoculation-timing cohort, we tested the effect of strain identity (five strains), nested within a binary treatment variable (inoculation versus control). To test inoculation effects at the end of the growing season, we conducted a single PERMANOVA combining the early- and late-season inoculation cohorts. We included the inoculation time (early- or late-season) and its interaction with strain identity and inoculation treatment to test whether inoculation effects on end-of-season community composition depended on inoculation timing.

To visualize these effects, we fitted a distance-based redundancy analysis (dbRDA) corresponding to each PERMANOVA. For simplicity, we used a combined six-level treatment variable representing the five strains and the control rather than nesting strain identity within the binary treatment. Similar to PERMANOVA, we partialled out host plant species and plot identity as conditioning variables. Ordinations were fitted with the ‘dbrda’ function in ‘vegan’, and the significance of the constrained terms was assessed using 9999 permutations. For each ordination, sample scores on the first two constrained axes were extracted with the ‘scores’ function. Group centroids were computed as the mean dbRDA1 and dbRDA2 score of all samples within each treatment, with standard error of the mean also computed for each axis. We tested differences in multivariate dispersion using Permutational Multivariate Analysis of Dispersion (PERMDISP) based on the same Aitchison distances. Two-week samples were analysed separately for the early- and late-inoculation cohorts, whereas end-of-season samples were pooled. Differences in distances to group centroids were assessed using 999 permutations, followed by Tukey-adjusted pairwise comparisons.

We tested the effect of inoculation on ASV richness using linear mixed-effects models with inoculation treatment, inoculation timing and strain identity as nested fixed effects and plot identity and host plant species as nested random effects. Similar to community composition analysis, we fitted separate models for the early- and late-season cohorts two weeks after inoculation, whereas the end of the season cohorts were analysed jointly. Richness analysis was implemented with ‘lmer’ function in ‘lme4’ package (v1.1.37) [42].

#### Relative contribution of inoculation to explaining bacterial community assembly

We assessed the variation in bacterial community composition explained by host plant species, plot identity, sampling time, inoculation treatment, strain identity and inoculation timing using a global PERMANOVA analysis. All samples were used in the analysis; host plant species, plot identity and sampling time were entered before the inoculation terms so that the contributions of inoculation treatment, timing and strain identity were estimated after accounting for spatial and temporal variation in community composition.

To assess the relative contribution of inoculation within each inoculation timing and sampling time, we conducted separate PERMANOVAs for early- and late-season inoculation at the two-week and end-of-season sampling time. For each model, we compared the variation explained by inoculation treatment and strain identity with that explained by host plant species and plot identity.

We used additional dbRDA ordinations to visualize how strain-associated compositional variation changed over sampling times within each inoculation cohort. Unlike the sampling-time-specific dbRDAs described above, these models combined the two sampling times and were fitted separately for the early- and late-inoculation cohorts. The interaction between the six-level treatment variable (five strains plus a control) and sampling time was used as the constrained term, while host plant species and plot identity were partialled out as conditioning variables.

## Results

### Inoculation increased the relative abundance of inoculated Methylobacterium taxa across strains and inoculation times

Sequencing of the *Methylobacterium*-specific *rpoB* marker gene revealed that inoculation greatly increased the relative abundance of ASVs matching the inoculated *Methylobacterium* strains compared to the non-inoculation controls (figure 2, figure S2). In the short term (two weeks post-inoculation), strains inoculated early in the growing season (May) dominated the *Methylobacterium* community with relative abundances ranging from 48 to 58%; the same strains inoculated at the late growing season (August) also established, although their relative abundance was lower (relative abundances from 28 to 51%). The relative abundance of inoculant-matched ASVs declined towards the end of the growing season, with a greater decline following early-than late-season inoculation, consistent with the longer interval between inoculation and end-of-season sampling. Nevertheless, at the end of season, inoculant-matched ASVs remained more abundant on inoculated leaves compared with control leaves for both early- and late-season inoculation treatments.

**Figure 2.**
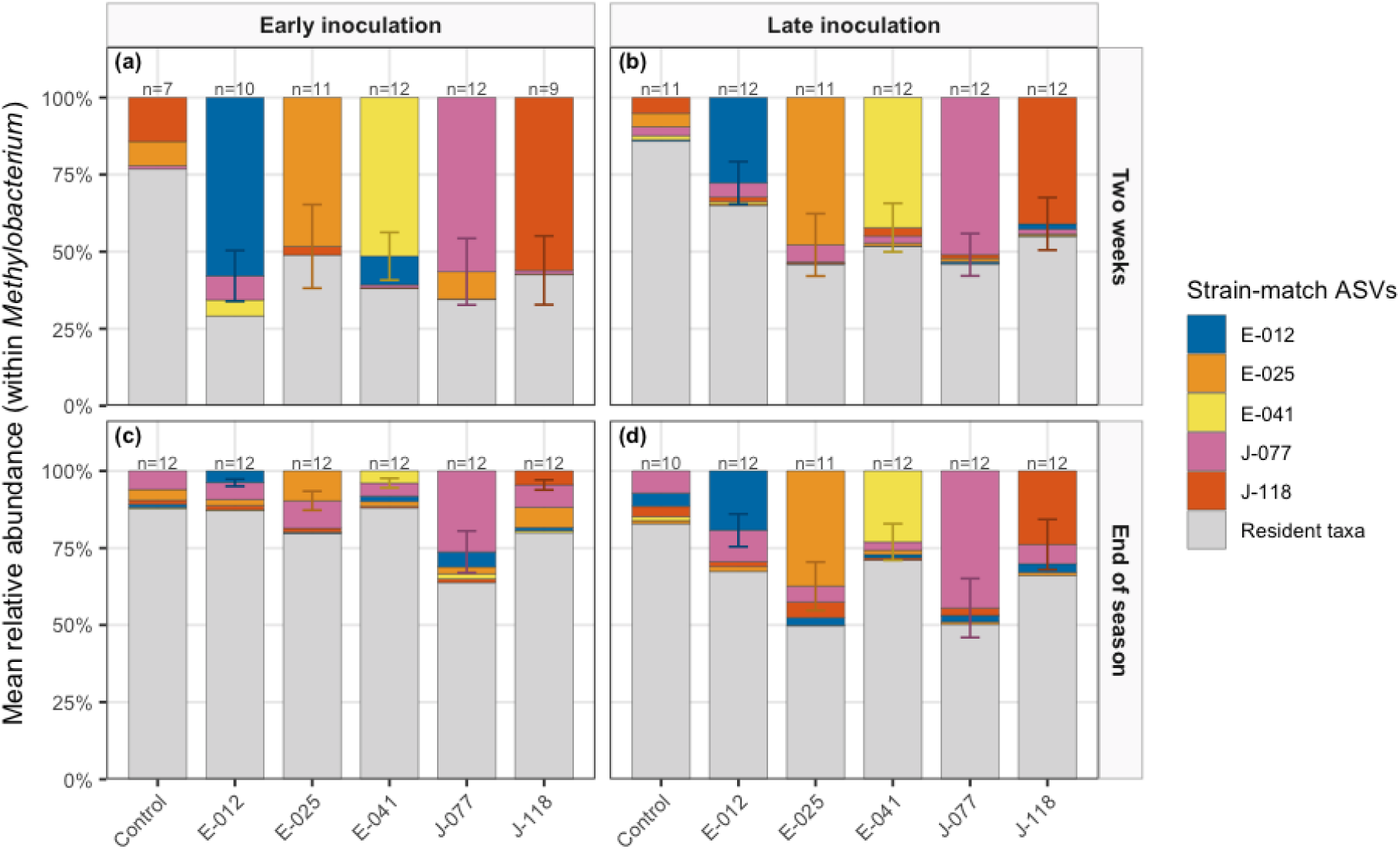
Relative abundance of inoculated strain-matched and resident *Methylobacterium* ASVs in phyllosphere communities, based on *rpoB* sequencing. Panel columns indicate inoculation timing (early-season inoculation, May; late-season inoculation, August) and panel rows are sampling stage (two weeks post-inoculation; end of season). The x-axis shows the treatment: an uninoculated control or one of the five inoculated strains. Stacked bars show the mean relative abundance of ASVs matching each inoculated strains, cloulors represent ASVs matching each reference strain, whereas gray shows resident taxa. Error bars are ±1 SE of the relative abundance for each inoculated strain (controls have none); n is the number of phyllosphere communities sampled. Labels above bars indicate the number of communities sampled.

Responses to inoculation differed among strains and depended on inoculation timing. E-012, a strain that was more abundant later in the growing season in natural communities (table S1), responded more strongly to early-season inoculation, reaching a relative abundance of 0.59 ± 0.08 (mean ± SE) in two weeks, than to late-season inoculation (reaching a relative abundance of 0.28 ± 0.07, figure 2). In contrast, J-118, which was naturally more abundant early in the growing season, reached a high relative abundance of 0.56 ± 0.11 shortly after inoculation, but declined substantially to 0.05 ± 0.02 at the end of the season (figure 2). Although the relative abundance of inoculated strains generally declined with time, late-season inoculated strains, especially J-077, remained abundant across communities at the end of the season.

### Inoculation of Methylobacterium strains enriched the abundance of non-Methylobacterium bacterial taxa

Sequencing of the 16S rRNA marker gene indicated that inoculation with *Methylobacterium* strains affected bacterial taxa beyond the *Methylobacterium* communities. Differential abundance analysis identified six 16S rRNA ASVs that were significantly enriched on leaves inoculated with *Methylobacterium* strains compared with non-inoculated controls, across both host plant species (figure 3). Three out of the six ASVs were assigned to *Methylobacterium*, whereas the remaining three were assigned to *Rhizorhabdus*, *Rhizobacter* and the *Oligoflexia* genus *0319-6G20*. Abundance enrichment of three *Methylobacterium* ASVs was most apparent two weeks after the inoculation, and diminished towards the end of the season, particularly following early-season inoculation. This result was consistent with our analysis of the *rpoB* marker, although here we were unable to make an exact match between 16S rRNA ASVs and the inoculated strains. In contrast, enrichment of the three non-*Methylobacterium* ASVs was associated with late-season inoculation: *Rhizobacter* was enriched two weeks after inoculation, whereas *Rhizorhabdus* and *0319-6G20* were enriched at the end of the season.

**Figure 3.**
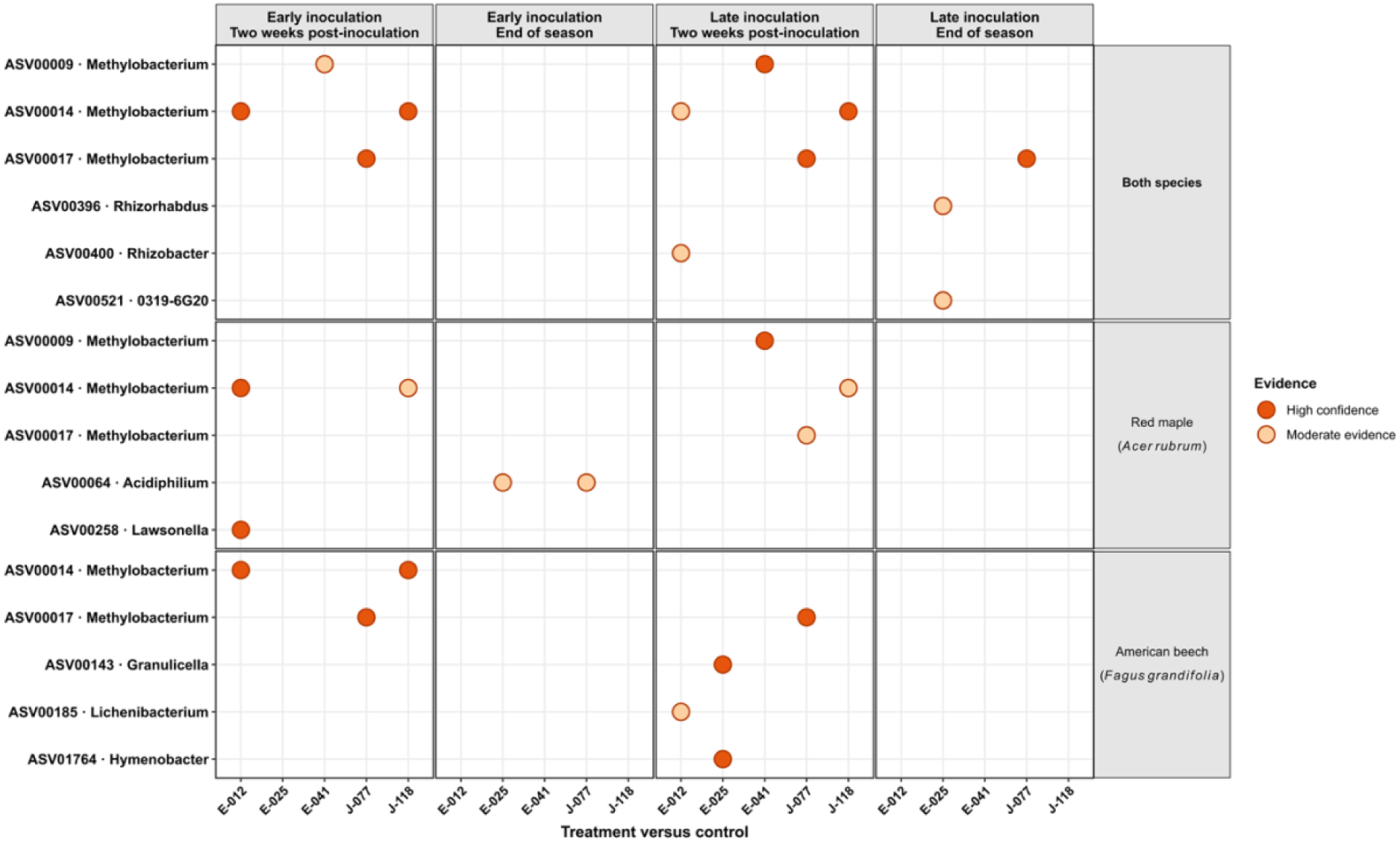
16S rRNA ASVs enriched on inoculated leaves versus uninoculated controls. Differentially abundant ASVs are shown for both host plant species combined, red maple, and American beech (rows), across inoculation timings and sampling stages (columns). The x-axis indicates the inoculated strain, and the y-axis indicates ASV identity and genus-level taxonomic assignment. Each point represents an ASV enriched in the corresponding inoculation treatment relative to the control. Five differential abundance methods were applied, and associations were retained when at least two methods agreed in the direction and statistical significance of the response. Dark orange points indicate high-confidence evidence (q < 0.05) while light orange points indicate moderate evidence (0.05 <= q < 0.1).

Since phyllosphere bacterial communities may differ among host tree species, we further conducted differential abundance analysis separately for each host tree species. Five enriched ASVs were identified for each host plant species, including three *Methylobacterium* ASVs and two non-*Methylobacterium* ASVs in red maple, and two *Methylobacterium* ASVs and three non-*Methylobacterium* ASVs in American beech (figure 3). Importantly, the enriched non-*Methylobacterium* ASVs differed between the two host plant species and among inoculation treatments. In red maple, ASVs assigned to *Lawsonella* and *Acidiphilium* were enriched with early-season inoculation. In American beech, ASVs assigned to *Granulicella*, *Lichenibacterium*, and *Hymenobacter* were enriched two weeks after late-season inoculation.

### Effects of inoculation on bacterial community composition depend on inoculation timing and strain identity

Two weeks after inoculation, we observed contrasting impacts of early- vs late-season inoculation on overall bacterial community composition compared with control leaves. Early-season inoculation led to strain-specific shifts in bacterial community composition (PERMANOVA strain identity nested within inoculation treatment; R2=6.3%, P=0.008). These shifts occurred in different directions relative to the controls (figure 4), and thus there was no consistent overall effect of inoculation on community composition (PERMANOVA inoculation-versus-control; R2=1.5%, P=0.21). In contrast, late-season inoculation did not significantly alter bacterial community composition (PERMANOVA inoculation-versus-control; R2=1.1%, P=0.65, strain identity nested within inoculation treatment; R2=4.8%, P=0.62, figure 4). Thus, strain identity had a stronger influence on the bacterial community composition when strains were introduced early rather than late in the growing season.

**Figure. 4.**
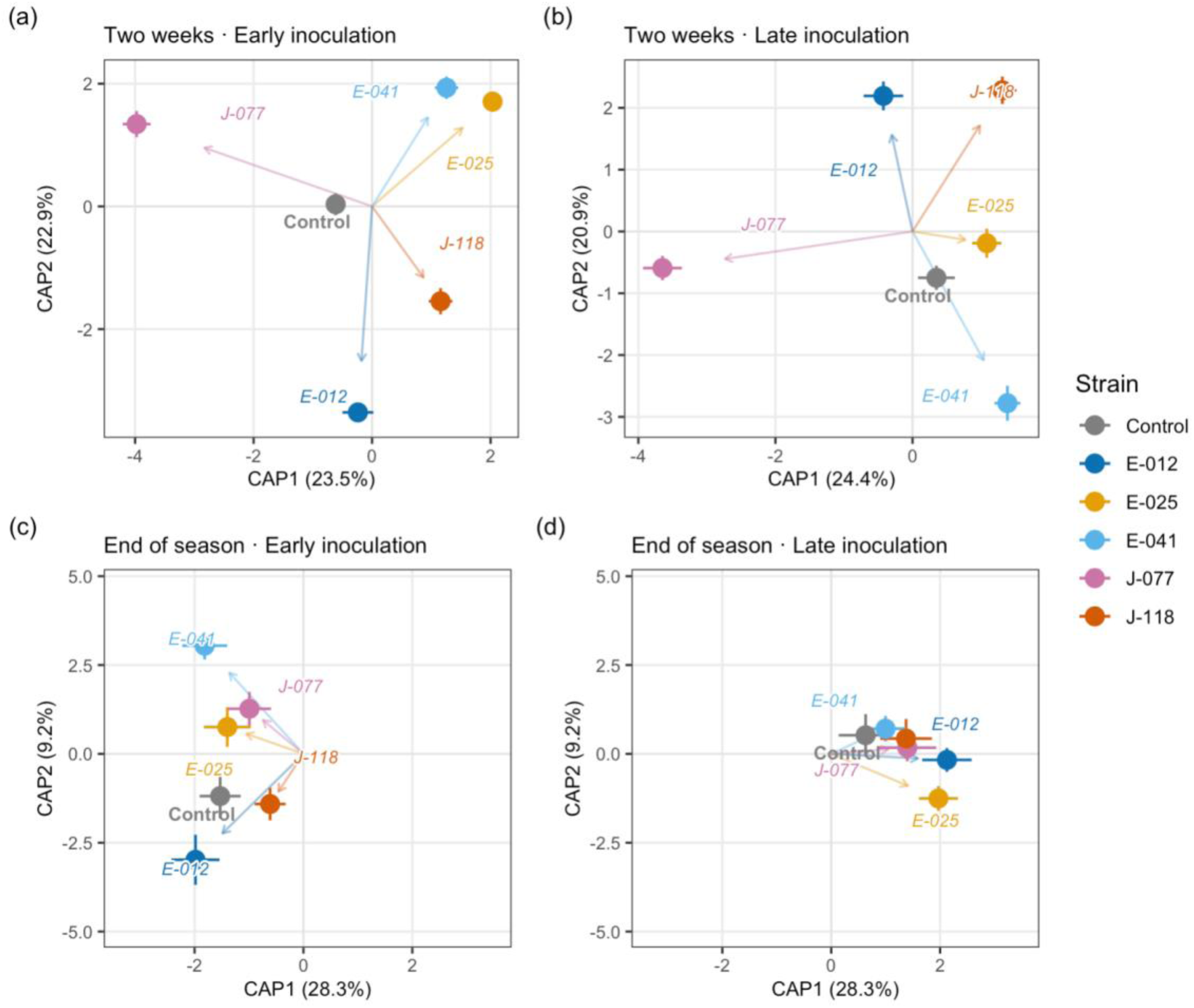
Effects of inoculation treatments on bacterial community composition (a, b) two weeks and (c, d) at the end of the growing season following (a, c) early-season and (b, d) late-season inoculation, using distance-based redundancy analysis (dbRDA) ordinations. Panels (a) and (b) were analyzed using separate dbRDA models as the samples were collected at different times of the season; inoculation treatment had a significant effect on community composition following early-season inoculation (P=0.033; a), but not following late-season inoculation (P=0.98; b). Panels (c) and (d) were analyzed from a combined model by incorporating an interaction term between inoculation treatment and inoculation timing (P=0.0129), thus sharing the same dbRDA axes. In all models, host plant species and plots were included as covariates. Points are group centroids, calculated as the mean dbRDA1 and dbRDA2 scores of all samples in each treatment; error bars show ±1 standard error on each axis. Arrows indicate the effect of each treatment, colors indicate the inoculated strains, and grey indicates uninoculated controls.

At the end of the growing season, we detected a significant interaction between inoculation treatment and inoculating timing in their effects on bacterial community composition (PERMANOVA inoculation-versus-control x early-versus-late-season inoculation interaction; R2=1.9%, P=0.001). Visualization using dbRDA showed that the same strain inoculated early versus late in the growing season resulted in distinct shifts in community composition (figure 4). Thus, the effect of inoculation timing remained detectable in bacterial community composition at the end of the growing season. Across all analyses, inoculation-related factors (treatment, inoculation time and strain identity) did not have detectable impacts on bacterial community ASV richness (P>0.05 in all models).

### Relative importance of inoculation effects on community composition diminished over time

Host plant species, plot, sampling time and inoculation timing all exhibited significant effects on phyllosphere bacterial community composition (PERMANOVA based on Aitchison distances; table 1). The total effects of inoculation-related factors, including inoculation treatment, inoculation timing and inoculated strain, explained 5.52% of variation in community composition. In comparison, environmental and temporal factors including host plant species, plot, season of sampling, and their interactions explained 10.46% of variation in community composition. Thus, inoculation played an important, although not dominant, role in shaping phyllosphere bacterial communities over the growing season.

**Table 1.** Global PERMANOVA of bacterial community composition based on Aitchison distances. The model included host plant species, site, their interaction, sampling time, inoculation treatment (control versus inoculated), inoculation timing (early versus late season, nested within inoculation treatment), and inoculated strain (nested within inoculation treatment). Model terms were fitted and tested sequentially in the order shown in the table using 999 unrestricted permutations. Df denotes the degrees of freedom, SS the sum of squares, pseudo F the PERMANOVA test statistic, and R^2^ the percentage of total variation in bacterial community composition explained by each term. Significance levels are indicated as ***P<0.001, *P<0.05, and ns, P**>**0.05.

| Variables | Df | SS | Pseudo<br>F | $R^2$<br>(%) | P-value |
| --- | --- | --- | --- | --- | --- |
| Host plant species | 1 | 3383.26 | 7.45 | 2.33 | < 0.001 *** |
| Plot | 1 | 6217.6 | 13.68 | 4.29 | < 0.001 *** |
| Host plant species × Plot | 1 | 1617.2 | 3.56 | 1.12 | < 0.001 *** |
| Sampling Time (Season) | 1 | 3948.8 | 8.69 | 2.72 | < 0.001 *** |
| Inoculation Treatment<br>(Control vs. Inoculated) | 1 | 517.1 | 1.14 | 0.36 | 0.168 ns |
| Inoculation Timing (Early vs. Late Season) | 1 | 3670.5 | 8.08 | 2.53 | < 0.001 *** |
| Inoculated Strain | 8 | 3805.3 | 1.05 | 2.63 | 0.283 ns |
| Residuals (Error) | 268 | 121779.2 |  | 84.0 |  |
| Total Variance | 282 | 144939.1 |  | 100 |  |

We further examined the relative contribution of inoculation-related and environmental factors at each inoculation timing and sampling time. Two weeks after the early-season inoculation, inoculation-related factors explained more variation in community composition (7.8%) than host plant species and plot combined (6.2%); however, the contribution of inoculation-related factors diminished over time, while that of host plant species and plots increased, explaining 5.7% and 16.8% of compositional variation at the end of the growing season, respectively. Following late-season inoculation, inoculation-related factors consistently explained less variation than host plant species and plot combined, both two weeks after inoculation and at the end of the season (5.6– 5.9% versus 18.6–19.6%). Visualization using dbRDA similarly showed that differences in community composition associated with inoculation became less pronounced over time, particularly following early-season inoculation (figure S3)

## Discussion

In this study, we inoculated bacterial strains onto tree leaves early or late in the growing season to quantify priority effects on phyllosphere community assembly. We found evidence for priority effects [8,19], namely that the timing of colonization of communities by inoculated *Methylobacterium* strains impacted community composition in different ways: early-season inoculation led to strain-specific shifts in community composition, while late-season inoculation had no effect [20,21]. The impacts of strain identity were strongest two weeks after early-season inoculation and weakened over time. At the end of the growing season, community composition differed between early- and late-inoculated leaves, indicating a longer-term influence of inoculation timing [13,16,21]. Our results thus supported our hypotheses that inoculation timing influences community assembly, and that the impacts of inoculation decrease over the growing season. While we detected priority effects on community assembly, the overall importance of inoculation-related effects on community assembly were smaller than the spatial, temporal and environmental impacts combined. Thus, microbiome engineering strategies seeking to exploit early-inoculation priority effects should also consider host and environmental context and continued immigration from surrounding microbial sources [13,16,21]. Our findings build on previous lab- and greenhouse-based studies [20,21] to provide evidence for the importance of priority effects over the growing season in a natural field-based setting [43].

Two weeks after inoculation, early- vs late-season inoculation led to contrasting patterns of community assembly. Early inoculation onto recently emerged leaves led to heterogeneous, strain-specific shifts in overall community composition, each strain-associated community type shifting in different ways compared to control leaves. Conversely, late season inoculation had no significant effects on community composition. These results are consistent with a model of phyllosphere community assembly where priority effects interact with the developmental state of host leaves and the successional state of communities on the leaves [18]. Furthermore, early colonizers might have the opportunity to alter the new leaf habitat more profoundly than late colonizers, or to engage in closer interspecific interactions with fewer established taxa, resulting in more diverse alternative trajectories for community succession [11]. In contrast, late inoculation may have limited impacts on community trajectories as mature leaves harbour abundant microbial populations and an already established community [22,23,43].

Strains showed important differences in terms of their ability to colonize and persist on leaves after inoculation [10], especially after early inoculation where there was significant heterogeneity in community composition. Strain J-077 was rare early in the season in natural communities (i.e. on uninoculated leaves); however, when inoculated early, it dominated *Methylobacterium* communities and remained among the most abundant strains on leaves throughout the season, even months after initial inoculation. This suggests that the low abundance of some strains in the early season may reflect delayed arrival; providing an earlier opportunity for establishment may allow them to dominate and gain a long-lasting priority advantage [44]. In contrast, strain J-118 was naturally more abundant early in the growing season and rare later in the growing season; when inoculated, this strain colonized the phyllosphere well initially, but it failed to persist, with its abundance declining greatly by the end of the season. Similar patterns of failure to persist after inoculation have been observed in another field inoculation study [43]. Therefore, late-season strains may benefit more from early inoculation, establishing large populations that persist throughout the season, while for early-season strains, the initial benefits from priority effects may eventually diminish as seasonal succession progress and environmental selection gains importance, consistent with their natural seasonal dynamics. Nevertheless, we observed that both early and late inoculation led to persistent and detectable changes in the composition of *Methylobacterium* communities, despite microbial dispersal, selection and other community assembly processes occurring in the natural environment.

Inoculation with *Methylobacterium* also enriched several non-*Methylobacterium* ASVs, suggesting indirect effects of inoculation on other bacteria, potentially mediated by microbial interactions. These effects differed between the two host plant species used in this study, consistent with previous studies showing important phyllosphere community variation among host plant species [14,17]. Although similar number of non-*Methylobacterium* ASVs were enriched by inoculation in the two host plant species, the identity of these enriched ASVs differed: two taxa belonging to *Lawsonella* and *Acidiphilium* genera were enriched on red maple after early inoculation with E-025 and E-012 or J-077, respectively, while other taxa belonging to *Granulicella* and *Lichenibacterium* were enriched on American beech after late inoculation with E-025 or E-012. These findings suggest that the indirect effect on non-*Methylobacterium* taxa depend on host plant species, inoculation timing and inoculated strains, highlighting the need to consider all three factors when developing microbiome engineering strategies [15,21]. Further investigating the role of host plant species in mediating priority effects on community assembly would be enlightening, just as would be evaluating the impact of spring inoculation onto evergreen conifer host plant species, whose leaves harbours bacteria throughout the year, forming communities distinct from deciduous trees [17,45].

While differential abundance analysis identified only a relatively small number of enriched ASVs following inoculation, multivariate analyses identified significant community-wide shifts in composition and dispersion. These community shifts could reflect changes in the abundance of many individual community members that did not pass the stringent statistical threshold used in differential abundance testing, or shifts in abundance that differed among inoculated strains, making it difficult to detect consistent responses among individual taxa to inoculation treatment [8]. Future studies that directly track the abundance of individual taxa through time will be required to identify which resident taxa respond to inoculation and how their response collectively generate shifts in overall community composition [44].

A central contribution of our study is that we quantified the relative importance of priority effect compared to host, spatial and seasonal processes under natural field conditions. Priority effects contributed substantially to community assembly, with effects comparable in magnitude to those of host, site and temporal factors. The contribution of inoculation-related factors declined over time, particularly following early-season inoculation, while host- and plot-related variation increased. This is consistent with evidence that environmental selection and species turnover can increasingly shape microbial communities during succession and reduce the influence of early colonists [46]. Microbial immigration from surrounding sources may further weaken the initial effect of inoculation [12]. These findings suggest that application of early inoculation may generate a strong priority effect in the short term but may not be sufficient to maintain persistent changes in community composition. Microbiome engineering strategies based on priority effects need to account for environmental filtering, microbial immigration and seasonal succession [7,47].

## Data availability

Sequencing reads including 16S rRNA gene and *rpoB* gene amplicon sequences are available in the NCBI Sequence Read Archive under BioProject accession PRJNA1471310. Whole-genome sequences of the inoculant strains are available under BioProject accession PRJNA730554.

Metadata and R scripts for data analysis are provided at https://github.com/skembel/phyllosphere-priority-effects-metadata-scripts.

## Acknowledgements

We thank Geneviève Bourret for assistance with library preparation and sequencing.

## Funding statement

This work was supported by a Natural Sciences and Engineering Research Council of Canada Discovery Grant (grant # RGPIN-2025-05608) to SWK.

## Notes

https://github.com/skembel/phyllosphere-priority-effects-metadata-scripts

https://www.ncbi.nlm.nih.gov/bioproject/PRJNA1471310

https://www.ncbi.nlm.nih.gov/bioproject/PRJNA730554

### Competing Interest Statement

The authors have declared no competing interest.

