## Supplementary Figures for "Priority effects influence bacterial community assembly on temperate deciduous tree leaves"

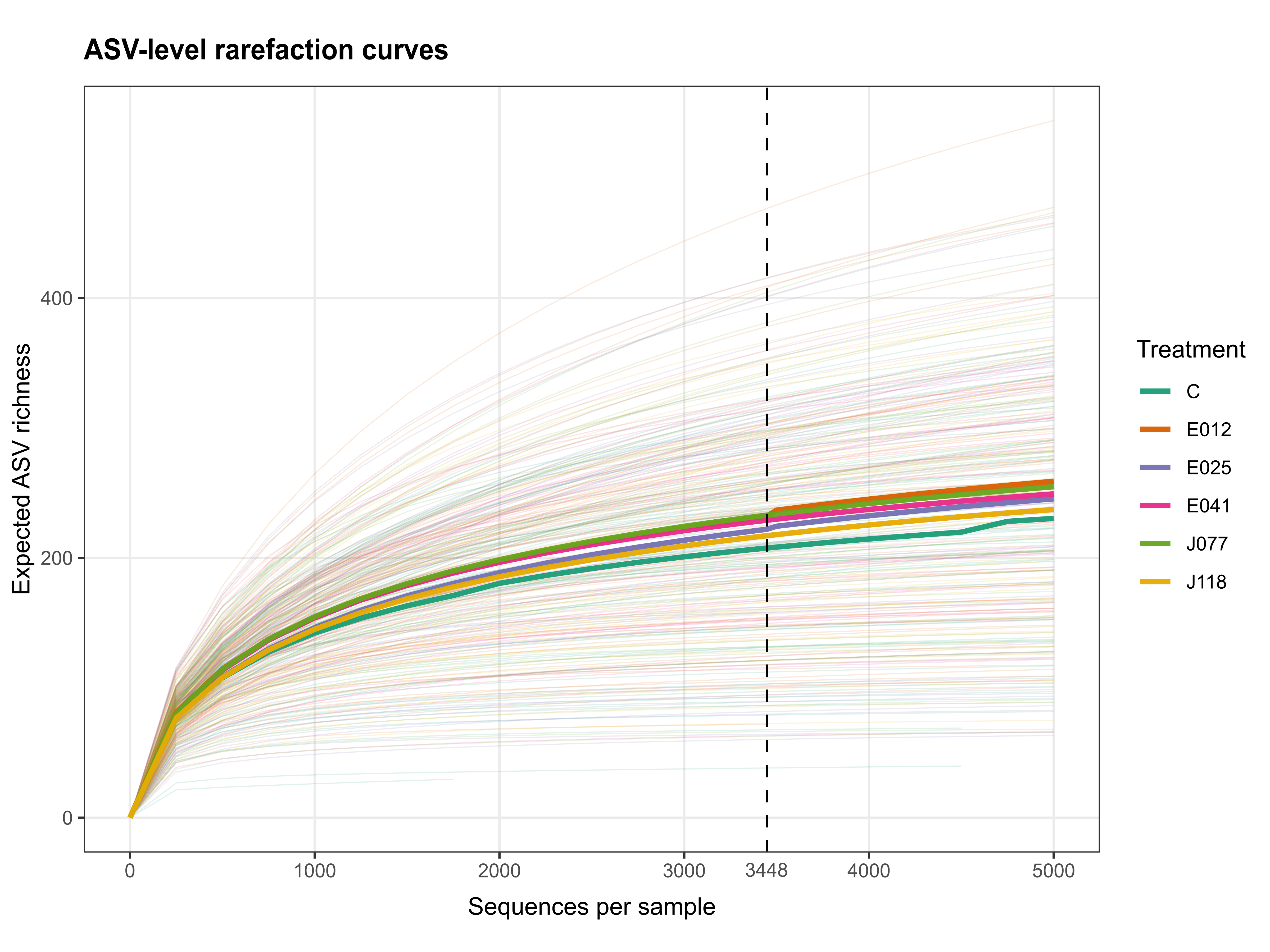

Figure S1. Rarefaction curve of number of ASVs as a function of number of sequences per sample for bacterial 16S rRNA data. Colours represent inoculation treatment; thin lines are individual samples; bold lines are treatment means. Means include samples with sufficient numbers of reads at each depth.

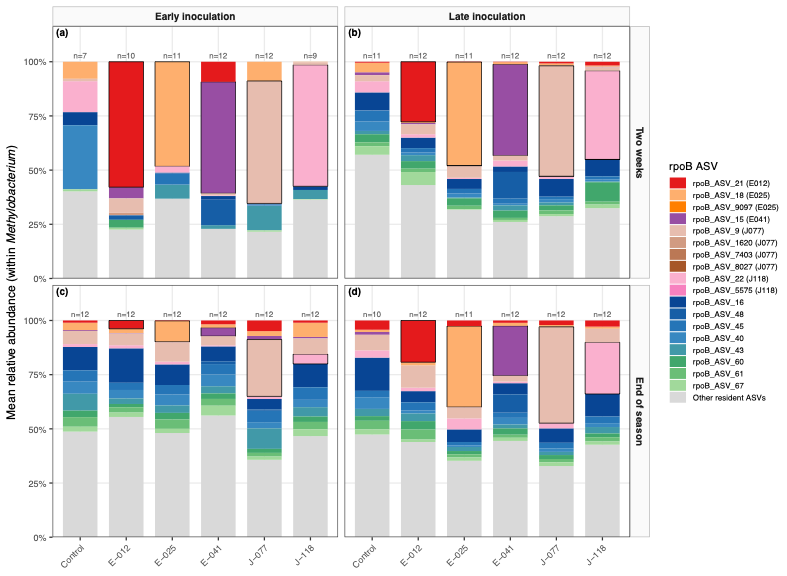

Figure S2. Relative abundance of *Methylobacterium* *rpoB* ASVs in phyllosphere communities. Panel columns represent inoculation timing (early-season inoculation in May; late-season inoculation in August) and panel rows show sampling time (two weeks post-inoculation or end of season). The x-axis shows the uninoculated control and the five inoculated strains. Stacked bars show the mean relative abundance of the 10 rpoB ASVs that uniquely matched the inoculated-strain references, the eight most abundant resident ASVs and all remaining resident ASVs pooled as “Other resident ASVs.” Black outlines identify ASVs matching the strain applied in each treatment. Labels above the bars indicate the number of phyllosphere communities included.

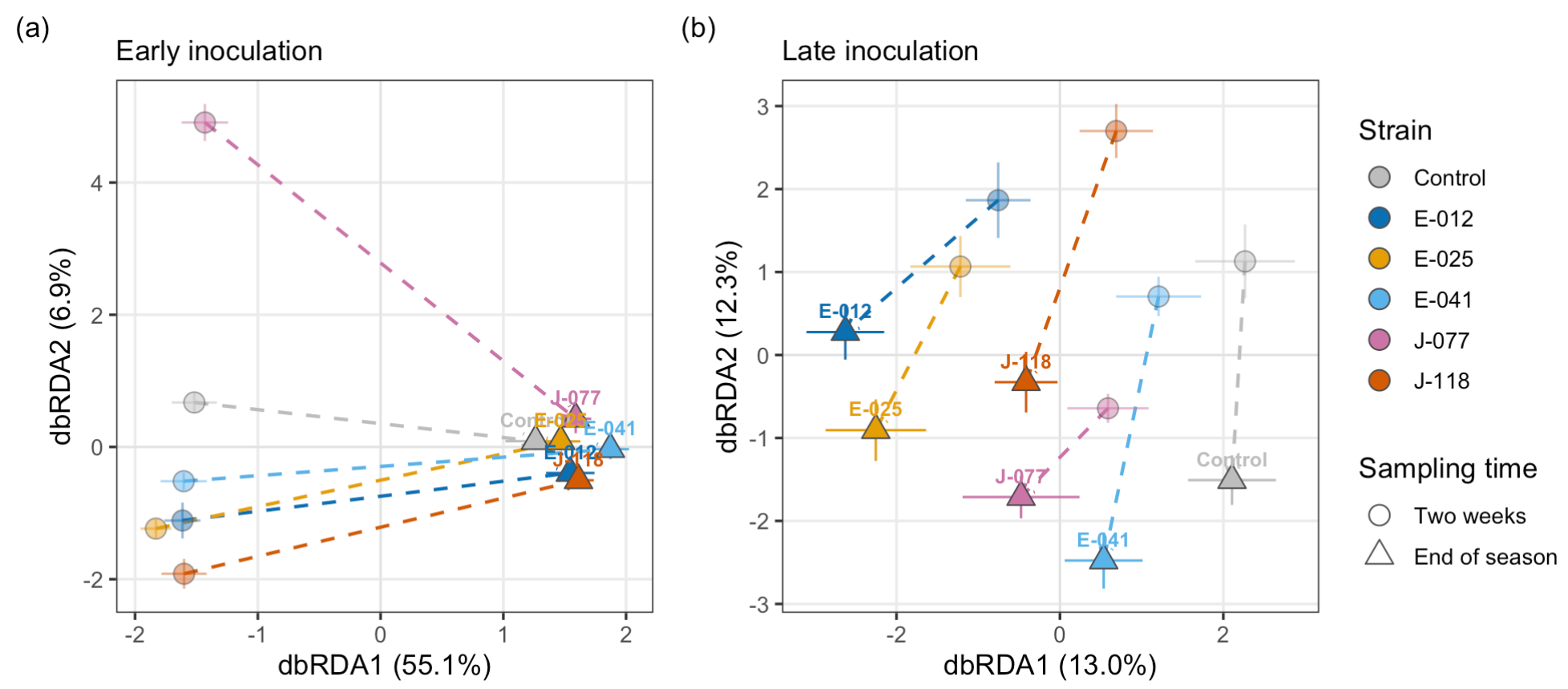

Figure S3. Temporal changes in bacterial community composition following (a) early- and (b) late-season inoculation. Points represent group centroids of samples inoculated with different strains (color) and collected at different stages (symbols), with error bars representing ±1 standard error on each axis. Dashed lines connect the two sampling stages for each treatment and indicate the direction of compositional change over time. Interaction between inoculation treatment and sampling time was used a constrained term, showing significant effects on community composition for early-season inoculation (P<0.001; a) but not for late-season inoculation (P=0.99; b), indicating inoculation effects changed with time following early inoculation.
